# Fungal communities (Mycobiome) as potential ecological indicators within confluence stretch of Ganges and Yamuna Rivers, India

**DOI:** 10.1101/848259

**Authors:** Rachel Samson, Vinay Rajput, Manan Shah, Rakeshkumar Yadav, Priyanka Sarode, Syed G. Dastager, Mahesh S. Dharne, Krishna Khairnar

## Abstract

River confluences are a hub of biodiversity with limited information with respect to the structure and the functions of the microbial communities. The ‘River Ganges’ is the national river of India, having manifold significance such as social, mythological, historic, geographic and agro-economic. It forms a sacred confluence (known as *Triveni Sangam*) with River Yamuna at Prayagraj, India. Recent reports indicated the presence of fecal coliform bacteria, an indicator recognized for water contamination in Ganges River leading to pollution. However, Fungi are also gaining attention as potential biological indicators of the trophic status of some rivers globally, but remain under-explored in terms of diversity, ecology and functional aspects. We performed whole long read, metagenome sequencing (MinION) of the sediment samples collected in December 2017 from confluence zone of Ganges and Yamuna Rivers spanning the pre-confluence, confluence and post-confluence zones. Mycobiome reads revealed a plethora of fungal communities, extending from saprophytes, endoparasites and edible fungi, to human pathogens, plant pathogens and toxin producers. The fungal genera recognized as bio-indicators of river pollution (*Aspergillus*, *Penicillium*) and eutrophication (*Kluveromyces*, *Lodderomyces*, and *Nakaseomyces*), were present in all samples. Functional gene analyses of myco-communities uncovered hits for neurodegenerative diseases and xenobiotic degradation potential, supporting bio-indicators of pollution. This study forms a foundational basis for understanding the impact of various anthropogenic activities on the mycobiome, as a bio-indicator of pollution across the confluence of Ganges and Yamuna Rivers and post-confluence of River Ganges, and could be useful in mitigating strategies for cleaning strategies of the Ganges River.

## 1. Introduction

Linking of the tributary/ tributaries to their main stream results in the formation of confluence (referred to as *Sangam* in Sanskrit language). Marked with an unusual geomorphology, aquatic ecology and variations in the distribution of the autochthonous communities, confluences house an array of biological diversity (Widder et al. 2014). However, our knowledge regarding the alterations in the fungal community structure as a result of river confluence is very limited.

Fungi are heterotrophic microbes having ubiquitous presence in the environment. It has been predicted that there are around 1.5 million fungal species of which only about 7% have been described (Mueller and Schmit 2007). Freshwater fungi have varied taxonomic and metabolic diversity and are fundamental in an efficient functioning of the food web dynamics of surface water eco-systems. In aquatic habitats, the presence of 3,000 fungal species and 138 non-fungal oomycetes has been estimated (Ittner et al. 2018). There are several studies that describe fungal diversity within various freshwater habitats such as ponds, rivers, streams and lakes (Bay and Kong 2004). Most of the freshwater fungi belong to the phyla *Ascomycetes*, *Basidiomycetes*, *Chytridiomycetes* and *Glomeromycetes* (Gulis et al. 2009). Some of the opportunistic pathogenic fungi in the water cause diseases in humans, aquatic plants and animals. Dispersion of fungal propagules is mediated by the flow of rivers and streams. The breakdown of the organic and aromatic wastes present in the water by these dispersed fungal communities, contribute to the nutrient recycling in the ecosystem (Luo et al. 2004). They are considered as essential components of the microbial loop, due to their dynamic involvement in the bio-transformation of xenobiotics, heavy metals and the decomposition of organic wastes of anthropic origin. In view of these aspects, fungi are regarded as important bio-indicators to assess the impact of anthropogenic intervention on the ecological and trophic state of a river or any aquatic body (Wang et al. 2015).

River Ganges is a dynamic riverine system of the world’s largest delta known as the Ganges-Brahmaputra-Meghna delta. It originates as River Bhagirathi from the Himalayan glacier- Gomukh and takes the name Ganges after the confluence of Bhagirathi River and Alaknanda River at Devprayag in Uttarakhand, India (Khairnar 2016). Ganges is the third largest river in the world in terms of the volume of water discharged. It spans approximately 2,525 km and drains about one-fourth of the Indian subcontinent. The religious, mythological, agricultural, industrial and economic importance of this river is marked since ancient times (Kumar 2017). Around 60% of the total waters of River Ganges is the result of its fifteen tributaries that converge with Ganges from its origin (Gomukh) to the point where it meets the Bay of Bengal (Zhang et al. 2018).

Amid all the tributaries of Ganges, River Yamuna is the longest tributary by length, making up 16% of the total water of Ganges joining Ganges at the *“Triveni Sangam”* in Prayagraj, India (Agarwal 2015). In Hindu tradition, *Triveni Sangam* is a sacred confluence of three rivers, namely- Ganges, Yamuna and the mythical *Sarasvati* River. Besides being the longest tributary, Yamuna River is among the most polluted rivers of India and upon its convergence with river Ganges, this pollution merges with that of the River Ganges (Misra and Misra 2010). Most of the studies pertaining to River Ganges in the past have emphasized on the deteriorating water quality and the alterations in the microbial diversity as an attribute of elevated anthropogenic activities (Farooquee et al. 2008; Bhardwaj et al. 2010). However, with the advent of metagenomic approaches and development in the sequencing technologies, it has become possible to comprehend microbial diversity in totality (Sanderson et al., 2018; Vey and Moreno-Hagelsieb, 2012a). The microbial (bacterial and archaeal) communities of Ganges with respect to human gut microbes and antibiotic resistance genes has been studied using Illumina sequencing technology (Zhang et al. 2018). In our previous study, we have profiled bacterial and archaeal communities across the confluence of River Ganges and River Yamuna (Samson et al. 2019). However, the fungal community structure and their functions across the confluence of River Ganges and River Yamuna remain unexplored. Therefore, to study the effect of allochthonous inputs on the myco-communities inhabiting these rivers is of the utmost importance. Targeting this facet, in the present study we have investigated autochthonous and the allochthonous mycobiome from the metagenome of Ganges River across its confluence with the Yamuna River, to gain insights into their taxonomic and functional diversity, using MinION sequencer, a paradigm shifting nanopore sequencing platform capable of real time sequencing of long reads of nucleic acids rapidly (Sanderson et al., 2018). Further, we have correlated the existing genera that mirror the human gut microbiome, domestic and urban pollution (sewage) to acquire clues about the impact of anthropogenic intervention on microbial load residing across the confluence of these rivers.

## 2. Materials and Methods

### 2.1 Sample collection and extraction of metagenomic DNA

A total of seven sediment samples were collected in triplicates (the left bank, center and the right bank), from each sampling site in the month of December 2017 (Fig.1). The samples were stored in ice and transported to the laboratory at 4°C for further microbiological analysis. Sediment samples from each location were pooled as 1 gram (gm.) from left + 1gm from middle + 1gm from right bank respectively, so as to encompass the total diversity (S1 Table). The total DNA was extracted using RNeasy PowerSoil Total RNA Isolation Kit (Qiagen, Germany) and RNeasy PowerSoil DNA Elution Kit (Qiagen, Germany) with certain modifications (S2 data). Each of the DNA samples were quantified by Nanodrop Lite Spectrophotometer (Thermo Fisher Scientific) and Qubit fluorometer (Thermo Fisher Scientific) using Qubit BR assay kit (Thermo Fisher Scientific, Q32853).

**Fig. 1:**
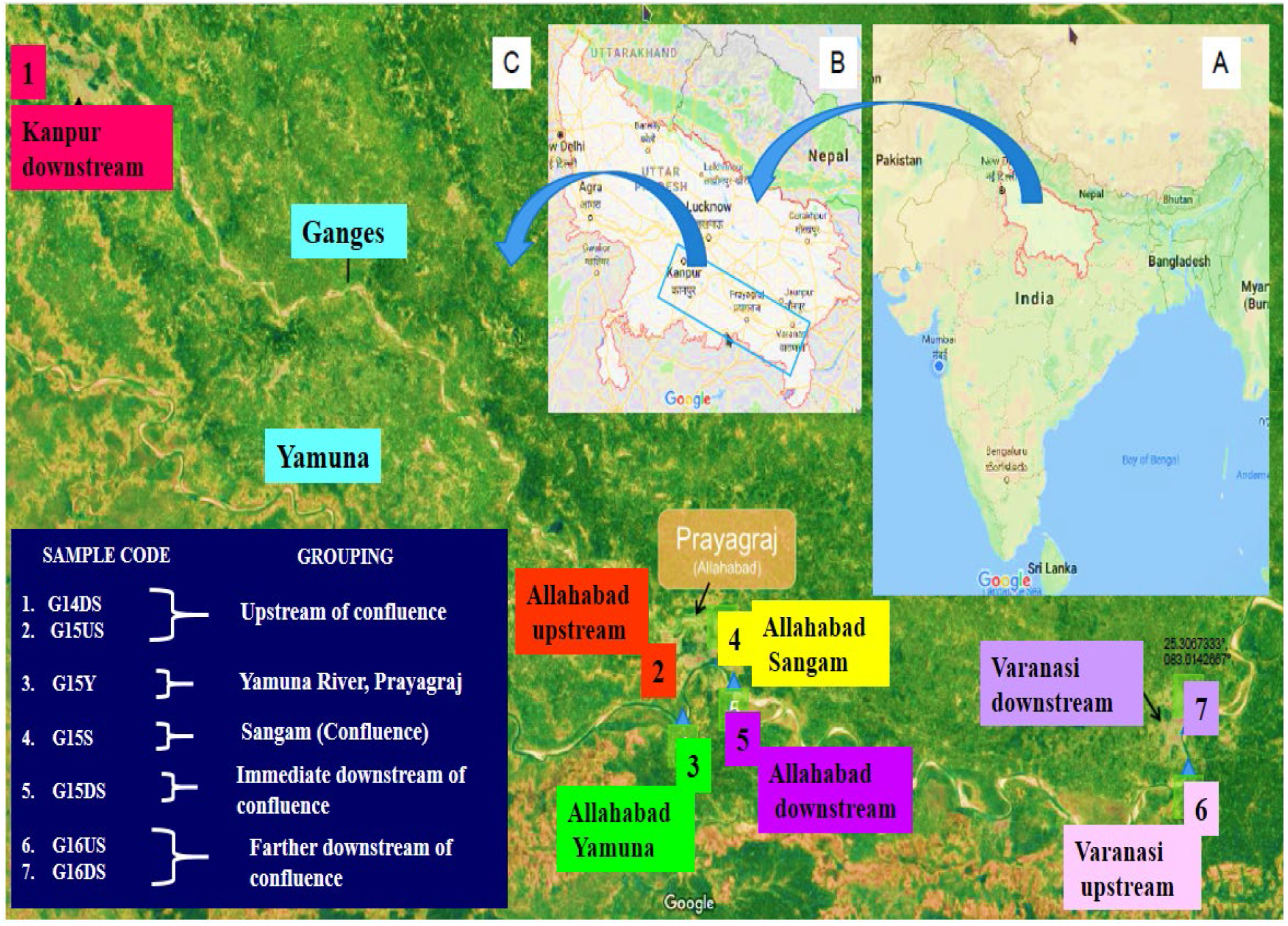
Details of the sampling locations and grouping of the samples investigated in this study (Samson et al. 2019).

### 2.2 High-throughput sequencing of the DNA samples

DNA extracted from the sediment samples of each location were sequenced by nanopore sequencing technology, using MinION sequencing platform. Preparation of the library was performed by the means of 1D Native barcoding genomic DNA (EXP-NBD103) and Ligation Sequencing Kit 1D (SQK-LSK108) protocol as suggested by the (Oxford Nanopore Technologies, Oxford, UK).

### 2.3 Pre-processing of the reads and Statistical analysis of the data

Albacore (v2.2.7) was used to base-call the reads. Poretools, a flexible toolkit for analysing the datasets produced by MinION nanopore sequencer was used for further Quality Check (QC) (Loman and Quinlan 2014). Error correction of the reads was achieved with Canu, to give good quality long reads (Koren et al. 2017). The analysis of taxonomic and functional diversity was carried out using MG-RAST (Metagenomic Rapid Annotation using Subsystem Technology) with an E value cutoff 1 × e^−5^ and sequence identity of 60% (Pozzo et al. 2018). Further, the taxonomic and functional analysis was based on the OTU table downloaded from the MG-RAST server. Analysis statistics of the assembled shotgun metagenome samples by MG RAST has been provided (Table Supplementary S3). First OTU distribution pattern was plotted in R followed by applying the Shapiro-Wilk normality test was also calculated to confirm the normal distribution of data. Parametric tests were applied to calculate the beta-diversity and alpha diversity.

Alpha-diversity plot was constructed using Simpson index and statistical test (t-Test/Anova) was also calculated based on the grouping of samples. Beta-Diversity was calculated using Microbiome Analyst to confirm whether a significant difference between two or more sampling groups exist (Anderson and Walsh 2013). A complete network plot of the functional profile was computed using Cytoscape v3.2.7 (Vey and Moreno-Hagelsieb 2012b) so as to gain insights into the functional potentials of the mycodiversity.

### 2.4 Availability of data

Metagenomics Rapid Annotation using Subsystem Technology (MG-RAST) server was used to allocate the taxonomic and functional diversity of fungal communities to all the seven metagenomic samples processed in this study. MinION sequence data for all of the metagenomic samples has been deposited at MGRAST server. MGRAST IDs assigned for the datasets are mgm4804931.3 (G14DS), mgm4804930.3 (G15US), mgm4804937.3 (G15Y), mgm4804936.3 (G15S), mgm4804939.3 (G15DS), mgm4804938.3 (G16US) and mgm4804941.3 (G16DS).

## 3. Results

### 3.1 Microbial diversity indices Assessment

Evaluation of diversity indices revealed the differences in complexity of myco-communities, typifying their miscellany across the confluence. The Shapiro-Wilk normality test/ W- test indicated normal distribution of data with p value of 0.04531 and a W statistic of 0.80444. Alpha diversity calculated using Simpson index which showed a relative increase in the fungal diversity for G15S indicating an increase in both the richness and the evenness of the myco-community at these locations as compared to the others. Another parametric statistical t-Test/ANOVA was applied to the sample groups based on the confluence, which showed significant difference in the fungal diversity of the sampling groups with a p-value of 0.46436, and F-value of 1.1187 (Fig. 2A). Beta diversity was calculated using PERMANOVA statistical test in MicrobiomeAnalyst using Bray-Curtis as the distance matrix. Analysis revealed significant differences between the groups with a p value <0.455, F value of 1.0137 and R-square value of 0.5034. From the analysis it was observed that the diversity of the confluence (G15S) and Yamuna river sample (G15Y) appeared to be similar. On the contrary, the diversity of samples in the immediate and farther downstream of confluence appeared to less diverse as compared to the confluence sample (G15S), while the G14US sample appeared to be most diverse in terms of fungal diversity (Fig. 2B).

**Fig. 2(A):**
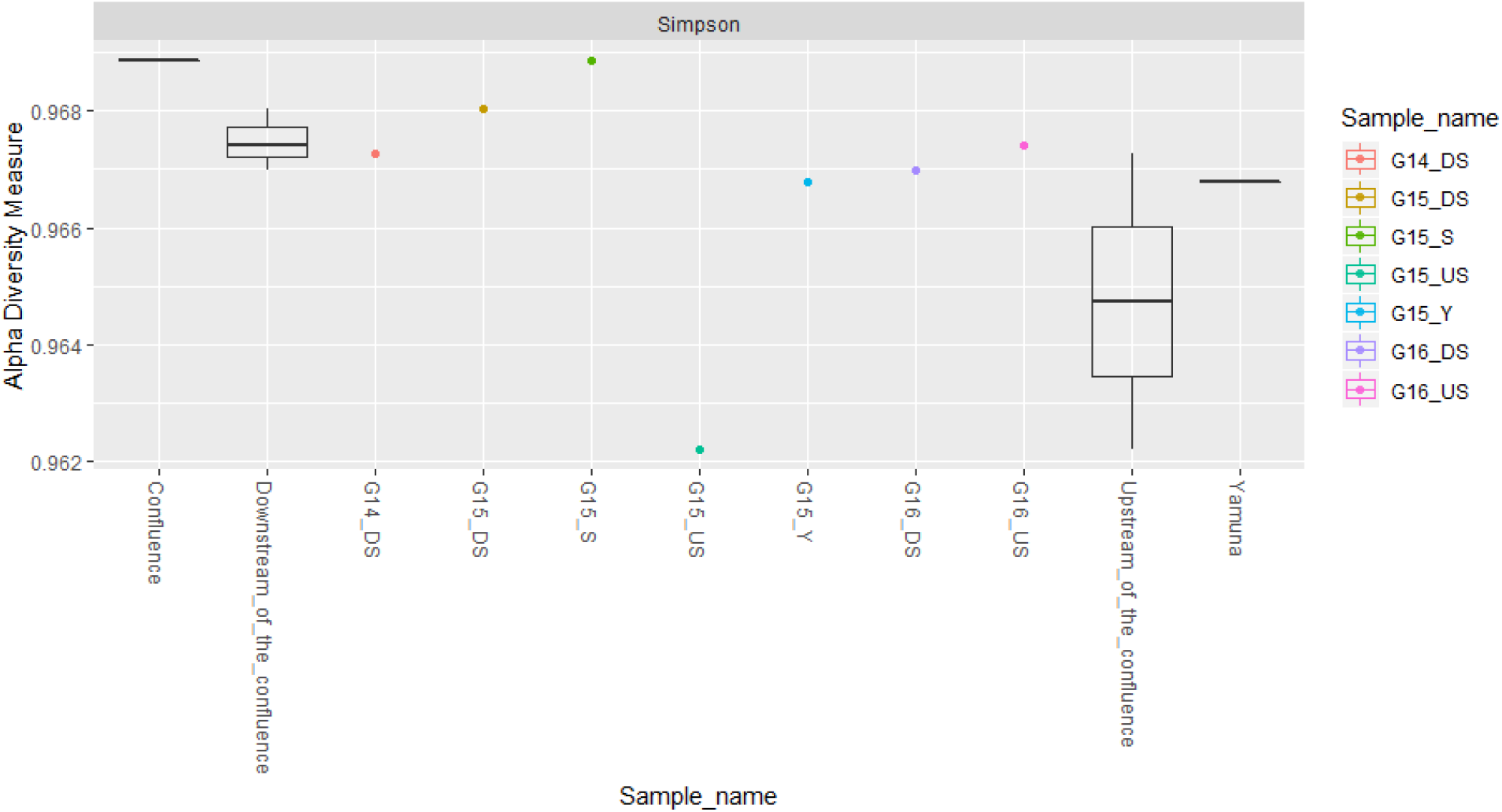
Simpson’s index of Alpha diversity measure describing the mycodiversity within each location under study.

**Fig. 2(B):**
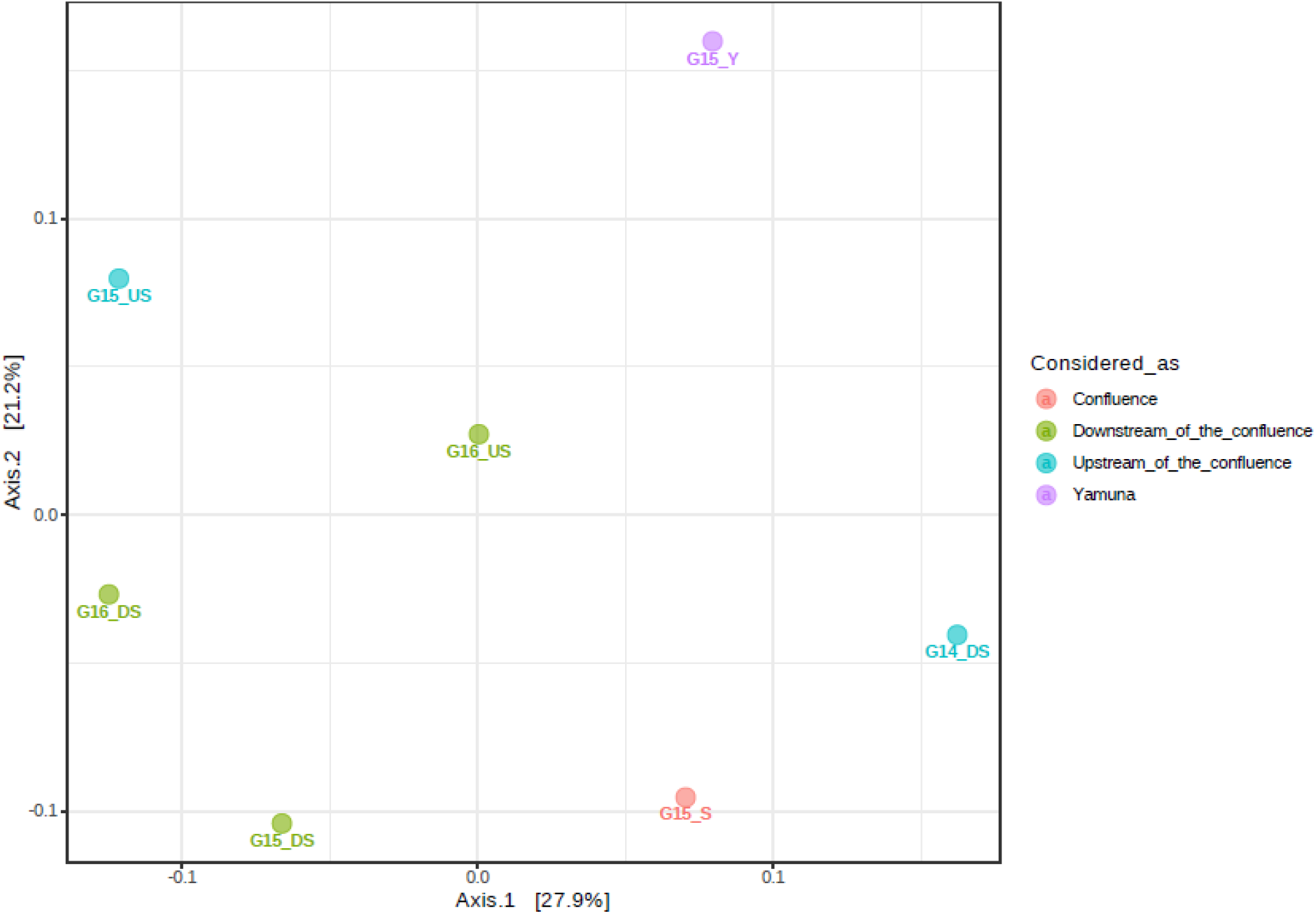
Principal Component Analysis (PCoA) plot using Bray-Curtis matrix showing Beta diversity of the mycobiome between the samples.

### 3.2 Fungal Diversity and Community Composition

MG-RAST analysis revealed 90-97% of the reads were assigned to the domain Bacteria, 1- 4% for Archaea and 2-3% for Eukarya. Bacterial and archeal reads were used in our previous study, to witness impact on the prokaryotic community in the wake of confluence with Yamuna River (Samson et al. 2019). Taxonomic profiling was performed using NCBI taxonomy data sets for all the samples which revealed 17 classified Eukaryotic phyla in each sample. Phylum *Ascomycota* predominated all the samples with a relative abundance of 15.28%, 13.37%, 14.94%, 15.91%, 14.14%, 12.97% and 14.12% corresponding to G14DS, G15US, G15Y, G15S, G15DS, G16US and G16DS respectively (Fig. 3A). Phylum *Basidiomycota* accounted for 2-4% in all the samples. However, *Chytridomycota* appeared to be absent in G15US, G16US and G16DS. In light of this, we further classified the fungal phyla so as to gain insights into the mycodiversity. At the family level, a higher relative abundance of family *Trichomaceae* (16-19%)*, Saccharomycetaceae* (11-13%)*, Nectriaceae* (5-7%) and *Debaryomycetaceae* (4-8%) dominated all the samples (Fig. 3B). Likewise, at the genus level, *Kluyveromyces* (13-20%), *Debaryomyces* (8- 14%)*, Giberella* (5-13%) and *Clavispora* (4-10%) were observed to be dominant at all the sampling sites. Notably, an upsurge in the relative abundance of *Rozella* (13%) and *Cladochytrium* (10%) was noted in G15US (Fig. 3C). On the basis of the taxonomic diversity at the genus level, we categorized fungi as bio-indicators of pollution due to anthropogenic activities (Fig. 4). Furthermore, classifying the genus level data of the myco-communities revealed a diverse ecology of fungi across the confluence (Table 1).

**Fig. 3:**
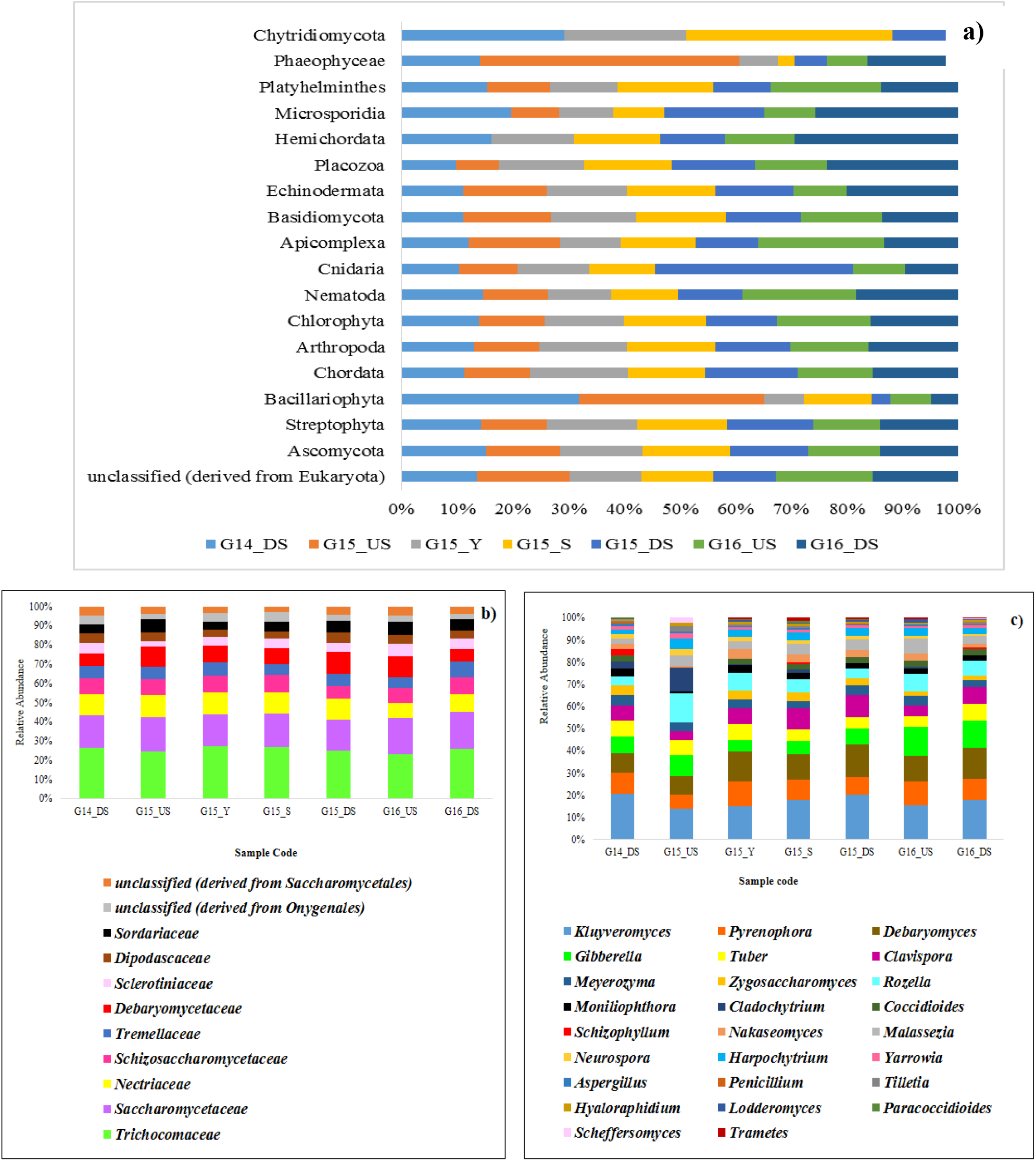
Myco-community composition **A)** Relative Phylum abundance of Eukaryotes; **B)** Relative Family abundance of Fungi (Top 10 families are shown); **C)** Relative Genera abundance of Fungi (Top 20 genera are shown).

**Fig. 4:**
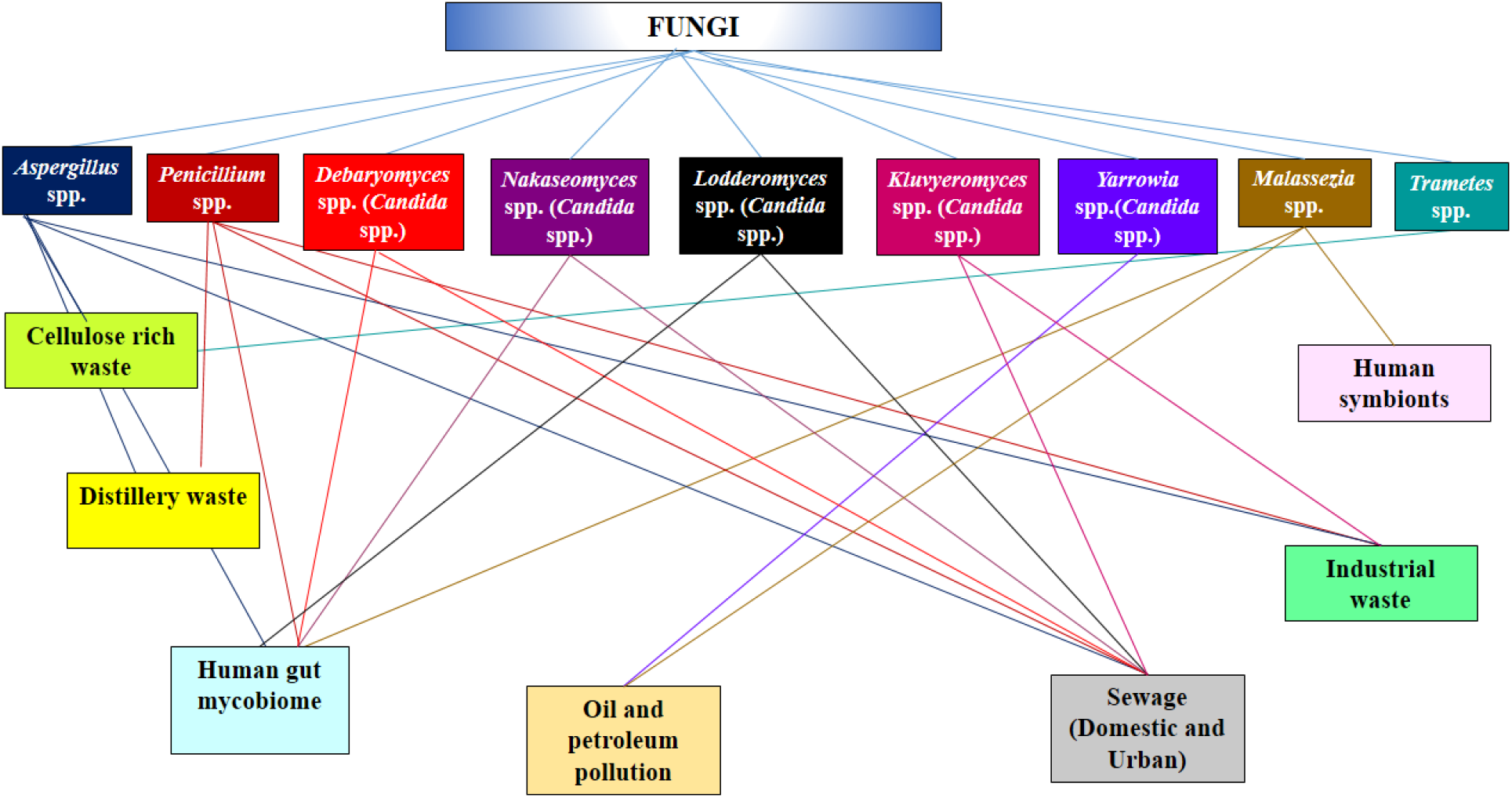
A network plot linking the bio-indicator group of fungi from all the sampling sites to its source.

**Table 1.** Classification of the fungal genera across the confluence of River Ganges and River Yamuna into different groups based on their ecology.

| Fungal Group | Genera | References |
| --- | --- | --- |
| Saprophytes | *Aspergillus*, *Penicillium*, *Trametes* | (Pitt 1994; Vélez and Díaz 2004) |
| Human Pathogens | *Clavispora*, *Paracoccidioides*,<br>*Coccidioides*, *Meyerozyma*,<br>*Nakaseomyces*, *Lodderomyces*<br>*Malassezia* | (Nosanchuk 2015; Köhler et al. 2015) |
| Plant Pathogens | *Gibberella*, *Pyrenophora*,<br>*Moniliophthora*, *Schizophyllum*,<br>*Tilletia*, *Cladochytrium* | (Doehlemann et al. 2017; Möller and Stukenbrock 2017; Termorshuizen 2017) |
| Edible Fungi<br>(Ectomycorrhiza) | *Tuber* | (Mello et al. 2010) |
| Unwalled Endoparasite | *Rozella* | (Gleason et al. 2012) |
| Killer Yeasts (Toxin<br>Producers) | *Kluyveromyces*, *Debaryomyces*,<br>*Zygosaccharomyces* | (Wang et al. 2008) |

### 3.3 In- Silico Functional Profile of the Mycobiome

KEGG (Kyoto encyclopedia of genes and genomes) database was used to understand the functional potentials of each sequence. KEGG Orthology (KO) numbers obtained from known reference hits and annotation of the metabolic pathways were assigned to each functional gene. In the first level of classification, metabolism accounted a major chunk (around 60%) of all the functions. The second level of classification involved in-depth analysis of the categories in level 1 to understand functional genes for amino acid metabolism, biosynthesis of secondary metabolites, carbohydrate metabolism, cofactors and vitamins metabolism, energy metabolism, glycan synthesis and metabolism, lipid metabolism, metabolism of terpenoids and polyketides, nucleotide metabolism and xenobiotic biodegradation and metabolism (Fig. 5A,B). Analysis of human disease revealed presence of neurodegenerative diseases such as Alzheimer’s disease (AD), Huntington’s disease (HD), Parkinson’s disease (PD) and Prions disease (PrD) at G15Y. It was observed that the fungal communities associated with HD are introduced into River Ganges only after its confluence with Yamuna River while the fungal taxa responsible for PD are confined to G15Y (Fig. 6). Further, the xenobiotic biodegradation aspect was analysed and annotated pathways for metabolism of xenobiotic compounds such as Chlorocyclohexane and Chlorobenzene and Benzoate were observed. Additionally, the potential for degradation of herbicides such as dioxin was observed only in G15Y and G15S (Fig. 7).

**Fig. 5:**
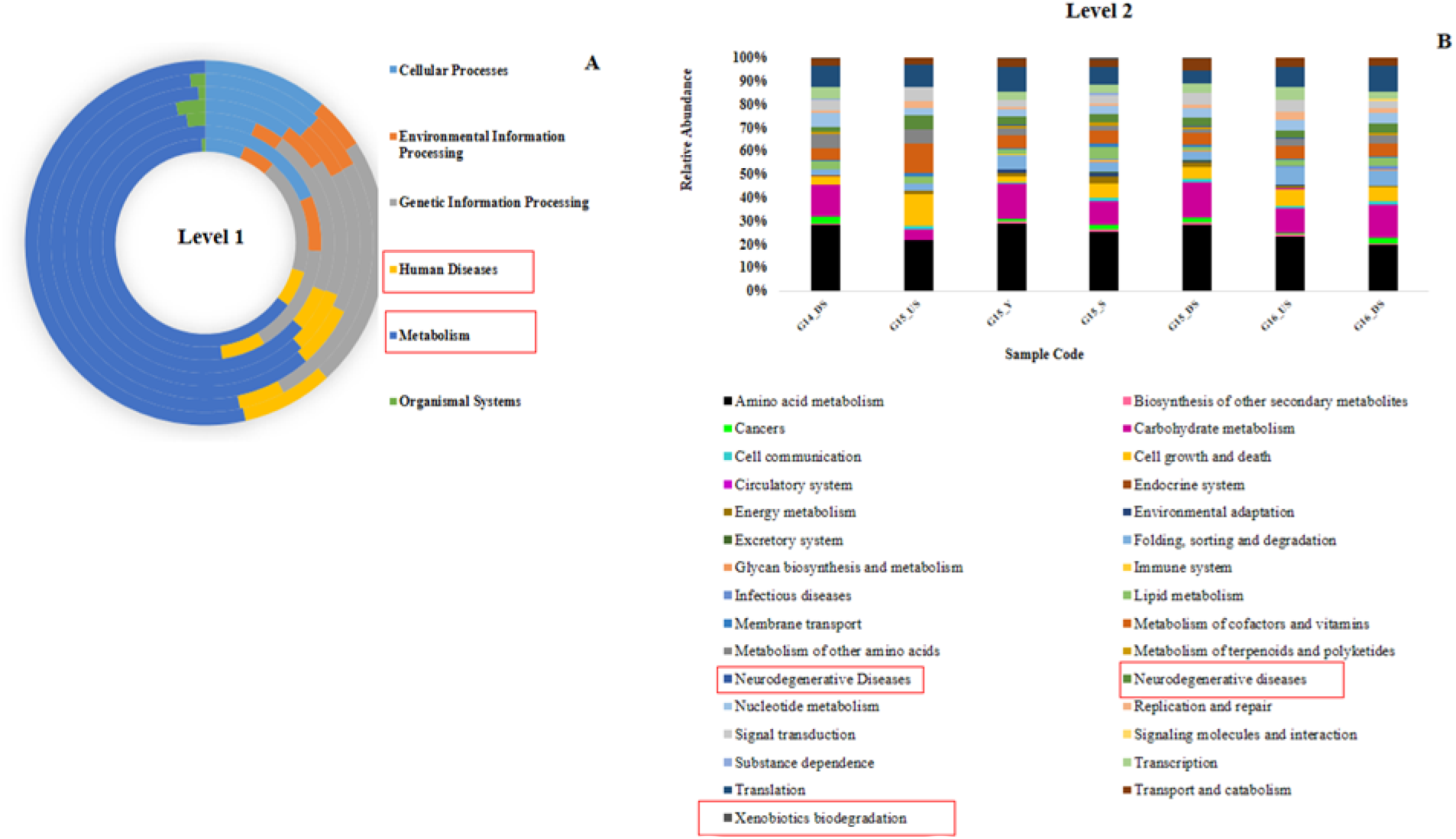
**(A)** Functional profile of the mycobiome by KEGG analysis at level 1. **(B)** Level 2 functional profile from KEGG analysis.

**Fig. 6:**
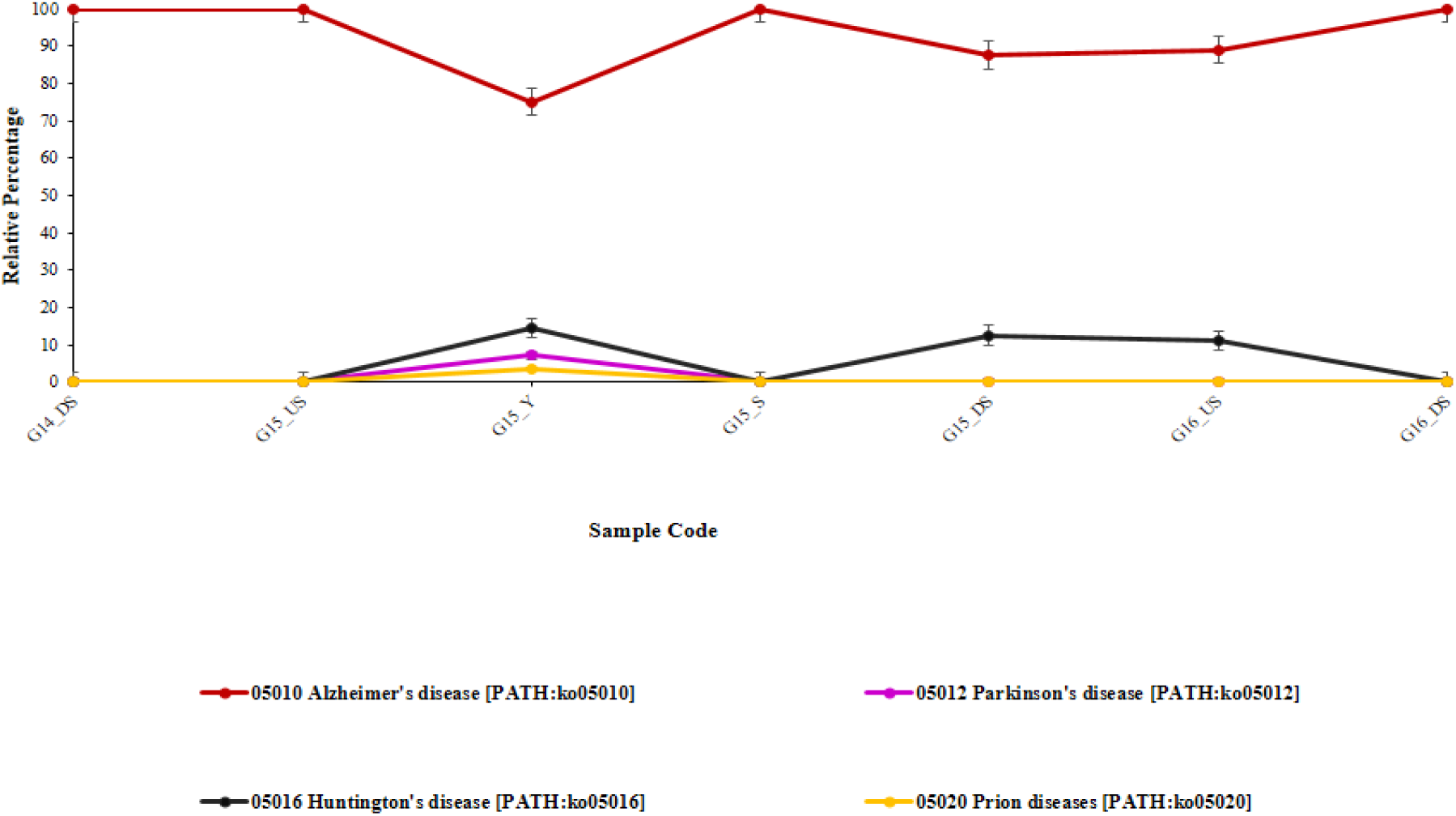
Graph showing hits against neurodegenerative diseases at different locations across the confluence of River Ganges and Yamuna.

**Fig. 7:**
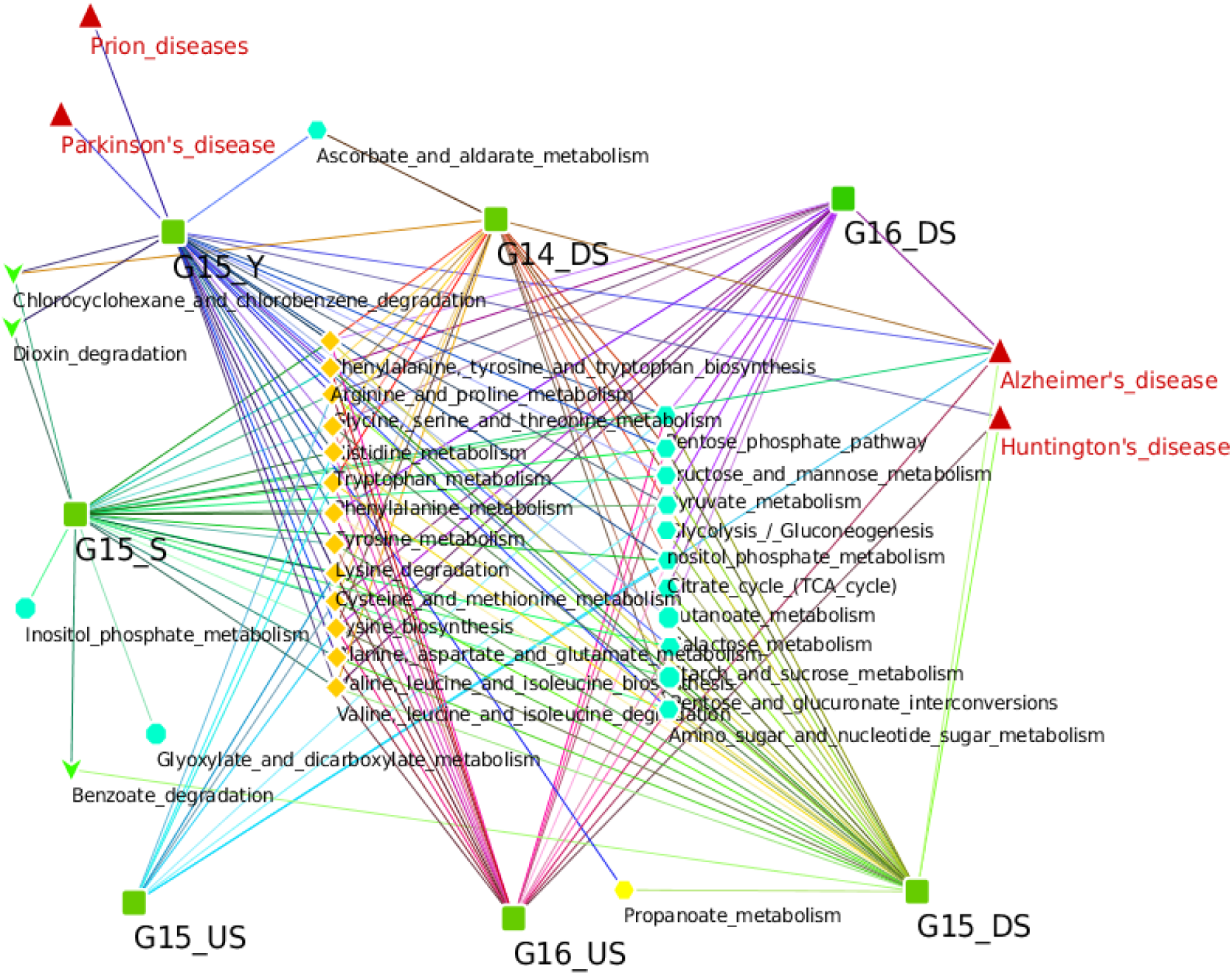
Network Plot depicting the entire functional gene(s) profiling of the Fungal Kingdom across the confluence of River Ganges and River Yamuna. Each location is indicated with a green box. Genes annotated for amino acid metabolism (yellow dots), Carbohydrate metabolism (blue dots). The connecting lines indicate functional genes shared in common by the locations, while gene(s) unique to a particular location are illustrated with arrows pointing outwards (red).

## Discussion

An exceptional biodiversity pattern is marked by the confluence of the River Ganga with River Yamuna. Till date, no studies have focused on the mycodiversity of these rivers across their convergence. Yamuna is one of the most polluted river in India and the organic load of pollution from this river adds up to River Ganges thereby amplifying the pollution levels. The presence of Fungi in polluted waters is considered beneficial for self-purification of the water (Biedunkiewicz, A. 2009). As majority of the microbial communities cannot be cultured under laboratory conditions, culture independent approach becomes the key to unlock the environmental test barrier (Chen et al. 2018). Targeting this facet, we have focused on revealing the fungal community structure and their associated functional genes using nanopore sequencing. Since sediment is less susceptible to ecological variation, it is more reliable as compared to the water counterpart (Li et al. 2016). Therefore, total metagenomic DNA was extracted from the sediment samples to comprehend the autochthonous (G14DS, G15US, G15S, G15DS, G16US, G16DS) and the allochthonous (G15Y) myco-community composition and the allied functional gene(s) content. MinION nanopore sequencer produces real-time data and encompass a broad range of applications in metagenomics and genome sequence assemblies (Brown et al., 2017; Sanderson et al., 2018).

Freshwater fungi are fundamental in functioning of the ecosystem because of their ability to decompose organic materials and recycle nutrients aiding to efficient biogeochemical cycling (Luo et al. 2004; Ittner et al. 2018). Besides this, they also play a key role in the nature as mutualists, parasites and pathogens (Schmit and Mueller 2007). In this study we present the first report, highlighting the mycodiversity of River Ganges and River Yamuna across their confluence. From the diversity measures it is evident that the pre-confluence of Ganges is comparatively less rich in terms of fungal diversity. However, an increase in fungal diversity at the confluence could be attributed to River Yamuna. Interestingly, the upsurge of myco-diversity remains confined only to the confluence, as the post confluence samples exhibit a decline in the fungal diversity, suggesting a transient influence of River Yamuna upon its confluence with River Ganges.

Analysis of the taxonomic diversity revealed dominance of phylum, *Ascomycota* in all the samples. This is comparable with the studies of other freshwater rivers like River Nile (Egypt), River Tietê (Brazil), River Kali (India) (Sudheep and Sridhar 2011; Abdel-Aziz 2016; Ortiz-Vera et al. 2018). *Ascomycota* is the largest, most diverse and ubiquitously distributed phylum of Fungi. The members of this phylum are crucial in the decay of autochthonous and allochthonous organic matter such as ligno-cellulosic waste from industrial and agricultural sectors (Gulis et al. 2009; Schoch et al. 2009). The decomposition of the organic matter by *Ascomycota* group is an important step in releasing inorganic nutrients to other microbial communities, thereby enhancing the nutrient cycling (Ma et al. 2013).

A higher relative abundance of family *Trichocomaceae* was observed in all the samples. Species belonging to this family are members of genus *Aspergillus* and *Penicillum*. Houbraken and Samson, 2011 have reported that genera of *Trichocomaceae* family have a diverse metabolic potential and secrete several pharmacologically important secondary metabolites such as penicillin, lovastatin, etc. Besides, the species of these genera have been reported to have a significant correlation with the fecal pollution (Arvanitidou et al. 2005). Another dominant group observed in this study was *Saccharomycetaceae* family, which includes phylogenetically circumscribed genera such as *Kluveromyces, Nakaseomyces* and *Zygosaccharomyces* from the “*Saccharomyces* complex” as descried by Van der Walt (van der Walt 1965). *Nakaseomyces* clade consist of pathogenic *Candida* species that are responsible for *Candia* infections (Angoulvant et al. 2015). *Kluveromyces, Lodderomyces* and *Nakaseomyces* are sexual stages of commonly occurring *Candida* spp. in the wastewater or the sewage sludge (Kutty and Philip 2008; Linder 2012). In addition to this, occurrence of these *Candida* spp. reflect the progressive eutrophic and pollution status across the confluence stretch, since they are well known bio-indicators of pollution (Wang et al. 2015). Furthermore, a higher relative abundance of uncommon opportunistic yeasts such as *Kluveromyces, Lodderomyces, Clavispora, Meyerozyma, Yarrowia* and *Malassezia*, previously isolated from bloodstream infections was also noted in this study (Lockhart et al. 2008; Taj-Aldeen et al. 2014; Cebeci Güler et al. 2017). Besides, other opportunistic group of fungal pathogens such as *Coccidioides, Paracoccidioides, Aspergillus, Penicillium* responsible for respiratory tract infections and onchomycosis were also detected (Vélez and Díaz 2004; Montes et al. 2015). Occurrence of these opportunistic fungi could be correlated with the epidemiological data of water-borne diseases acquired by the bathers after the mass bathing event of *Kumbh mela* taking place every 6 or 12 years at the confluence of River Ganges and River Yamuna (Tyagi et al. 2013), while the existence of bio-indicator species of pollution could be correlated with the massive anthropic inputs in these rivers (Misra and Misra 2010; Zhang et al. 2018).

Another important observation made in this study was an upsurge in the relative abundance of *Rozella*, which belongs to *Cryptomycota*- a recently discovered phylum of early-diverging fungi*. Rozella* is a biotroph found in fresh as well as marine waters and are involved in modifying the dynamics of the aquatic food chain positively due to proficient transferal of carbon and energy from the primary consumers (their hosts) to the secondary and tertiary consumers (Gleason et al. 2012). Further, appearance of genera such as *Harpochytrium* and *Hyaloraphidium* belonging to *Chytridomycota* was noted. These genera play an important role in decomposition of the organic particulate matter and conversion of inorganic compounds to organic compound increasing their bioavailability to other microbes of the aquatic food web (Gleason et al. 2008). Alterations in the myco-community structure as a result of confluence were not observed at the phylum, family and the genus level. Persistence of an unchanging pattern of mycodiversity pre-confluence and post confluence suggest a transient influence of River Yamuna upon its confluence with River Ganges at present.

Discovery of novel genes that confer diverse metabolic potentials to the microbes could be achieved through the metagenomic study. Progressive rise in the pollution levels of Ganges and Yamuna rivers due to added anthropogenic, industrial, agricultural and domestic wastes, makes it crucial to investigate the functional potentials of myco-diversity. Aiming there-to, we have focused primarily on the genes involved in carbohydrate and amino-acid metabolism along with those contributing to neurodegenerative diseases and metabolism of xenobiotic compounds. The functional annotation of the metagenomes was done through MG-RAST using KEGG analysis. Several metabolic functions like biosynthesis of secondary metabolites and amino acid, carbohydrate, energy, lipid, cofactors and vitamins, terpenoids and polyketides, nucleotide and xenobiotic biodegradation and metabolism were observed in the analysis. There are several reports on occurrence of fungi in different regions of brain tissues and blood vessels, recovered from the patients suffering Alzheimer’s disease and Parkinson’s disease (Pisa et al. 2015; Alonso et al. 2017, 2018). Addressing this aspect, we emphasized our analysis towards the functional profile of the autochthonous and the allochthonous fungal communities. The analysis revealed hits for AD, HD, PD and PrD which fall under the category of neurodegenerative diseases of human diseases. A probable microbial etiology for AD and PD has been proposed by a number of researchers (Alonso et al. 2015; Pisa et al. 2015). Detection of hits against AD and PD in this study could be correlated with the anthropogenic and ritualistic activities, floatation of decaying corpses, immersing of dead body into River Ganges prior to cremation, etc. (Sharma 1973; Kedzior 2014). The breached integument would probably serve as a gateway, releasing the body fluids from the affected patients into the river. Furthermore, presence of hits against prions disease exclusively in Yamuna river sample was noteworthy. Prions are infectious, misfolded proteins which characterize several amyloid related neurodegenerative diseases (Wickner et al. 2015). Several Yeasts like *Kluveromyces, Saccharomyces, Candida* and *Pichia* have been reported with the ability to form prion proteins (Baudin-Baillieu et al. 2003; Wickner et al. 2015; Friedmann et al. 2016). Introduction of such misfolded proteins into the river water of Yamuna could be a result of cell death or apoptosis of the fungus *Kluveromyces*.

Further we explored the fungal communities for their abilities to degrade xenobiotic compounds. Genes involved in degradation of xenobiotic compounds accounted about 0.5-0.9%. Classification of this aspect in detail revealed highest percentage of reads assigned to degradation of chlorocyclohexane and chlorobenzene, followed by dioxin and benzoate degradation. Chlorocyclohexane and Dioxins have a recalcitrant nature environment and therefore are amongst the principal xenobiotic compounds (Singh and Kuhad 1999; Kumar et al. 2012). Release of industrial or agricultural wastes carrying these compounds into these rivers has resulted in the pollution of the riverine waters and an adverse effect on the microbes residing therein. Here, we report for the first time, the presence of hydrocarbon assimilating genera such as *Clavispora*, *Debaryomyces*, *Lodderomyces* and *Yarrowia* across the confluence of River Ganges with River Yamuna. White-rot fungus like *Trametes* are well-known for their lignolytic properties. However, the extracellular peroxidase enzyme of this fungus is non-specific and has been associated with degradation of xenobiotics like chlorocyclohexane and chlorobenzene (Diez 2010; Meuser 2013). Saprophytic fungi like *Aspergillus* and *Pencillum* have been reported to degrade chlorocyclohexane (Maqbool et al. 2016). Besides, *Trametes, Aspergillus* and *Penicillium* have been reported for their ability to degrade dioxin (Chang 2008; Deshmukh et al. 2016; Pathak and Navneet 2017). Interestingly, degradation of dioxin appeared to be limited to the G15Y and G15S only and were not carried downstream of the confluence, suggesting momentary effect of Yamuna River only on the confluence. To the best of our knowledge, there are no reports highlighting the potentials of myco-communities in the degradation of these xenobiotic compounds across the confluence of River Ganges and River Yamuna.

## Conclusion

Our findings highlighted the taxonomy and functional potentials of the mycobiome across the confluence of River Ganges and River Yamuna using high throughput nanopore sequencing. The functional potentials of the fungal communities of Ganges River are transiently affected by the confluence of River Yamuna at present. Nevertheless, the taxonomic diversity of the mycobiome of River Ganges remains dismally affected by the confluence. Further experimental work needs to be carried out to understand and draw inferences on the enigmatic effects of anthropic intervention on the fungal diversity across the confluence of River Ganges with River Yamuna.

## Authors Contribution

KK, MSD, SGD designed and supervised study. MS, VR, RK, KK involved in sampling. RS, PS, RKY processed samples, library preparation and generated data. VR, MS performed data analysis. RS, MSD, VR, SGD, KK drafted the manuscript. All authors carefully read and approved the manuscript.

## Conflicts of interest

Authors declare no competing interests.

## Acknowledgements

Authors are thankful to Ministry of Jal Shakti, Department of Water Resources, River Development, and Ganga Rejuvenation, National Mission for Clean Ganga (NMCG), Government of India, New Delhi, India for financial assistance (GKC-01/2016-17, 212, NMCG- Research), Directors of CSIR-NCL, and CSIR-NEERI for infrastructure and support. RS and PS would like to acknowledge NMCG for fellowship. RKY acknowledges University Grants Commission (UGC), New Delhi, India and Academy of Scientific and Innovative Research (AcSIR), New Delhi India. The manuscript has been checked for plagiarism using iThenticate Software and assigned KRC No.: CSIR-NEERI/KRC/2019/MARCH/EVC/1.

