## Supplementary information for "Fungal communities (Mycobiome) as potential ecological indicators within confluence stretch of Ganges and Yamuna Rivers, India"

**Supplementary S1.** Details of the sampling locations and year of the samples processed in this study.

| Sample code | Location | Latitude | Longitude | Sampling year |
|---|---|---|---|---|
| G14DS | Kanpur downstream | $26^{0}$37.9005' N | $80^{0}$49.1344' E | Dec'17 |
| G15US | Allahabad upstream | $25^{0}$51.6161' N | $81^{0}$68.5440' E | Dec'17 |
| G15(S) | Allahabad Sangam | $25^{0}$42.3628' N | $81^{0}$88.8116' E | Dec'17 |
| G15(Y) | Allahabad Yamuna | $25^{0}$32.0510' N | $81^{0}$78.9160' E | Dec'17 |
| G15DS | Allahabad Downstream | $25^{0}$39.3014' N | $81^{0}$91.2330' E | Dec'17 |
| G16US | Varanasi upstream | $25^{0}$13.778'N | $83^{0}$01.513'E | Dec'17 |
| G16DS | Varanasi downstream | $25^{0}$18.404'N | $83^{0}$0.856'E | Dec'17 |

**Supplementary S2.** Modified protocol for extraction of Metagenomic DNA

Total DNA was extracted from the samples using RNeasy PowerSoil Total RNA Isolation Kit (Qiagen) and RNeasy PowerSoil DNA Elution Kit (Qiagen) with certain modifications. In brief, 2gm of the sediment sample was added to 15 mL bead tube with subsequent addition of 2.5 mL and 0.25mL of bead solution, solution SR1 respectively. To the above mixture in the bead tube, 3.5 mL of Phenol: Chloroform: Isoamylalcohol solution (pH 6.5) was and vortexed for 15 mins until the biphasic layer disappeared. This was followed by centrifuging the tube at 10,000 rpm for 10 mins at room temperature.

The upper aqueous layer was then transferred to a new 15 mL tube followed by addition of 1.5 mL of solution SR3. The resulting solution was then vortexed and incubated at 4°C for 10 mins. Post incubation, the samples was centrifuged for 10 mins at 9000 rpm and the supernatant was transferred into a new 15 mL collection tube, followed by addition of solution of SR4 and subsequent incubation at -20°C for 30 mins.

Further, this mixture was centrifuged at 9000 rpm for 30 mins. The supernatant obtained at this step was discarded and the pellet for each samples were re-suspended completely in 1mL of solution SR5. This suspension was then subjected to RNA capture column and the nucleic acids (DNA and RNA) were eluted using appropriate eluents like solution SR8 and SR6 respectively.

To each of the eluate, 1mL of SR4 solution was added and then incubated at -20°C for 10 mins. Later, the eluates were centrifuged for 15 mins at 13,000 rpm so as to pellet out their respective nucleic acids. The pellet obtained for each nucleic acid (DNA and RNA) was resuspended in 100µL of SR7 and stored at -20°C. The RNA was stored at -80°C

The DNA quality and concentration for each sample was assessed by Nanodrop Lite Spectrophotometer (Thermo Fisher Scientific) and Qubit fluorometer (Thermo Fisher Scientific) quantification using qubit BR assay kit (Thermo Fisher Scientific, Q32853).

**Supplementary S3**. R script for analysis

**Supplementary S4.** Rarefaction plots of the mycobiome

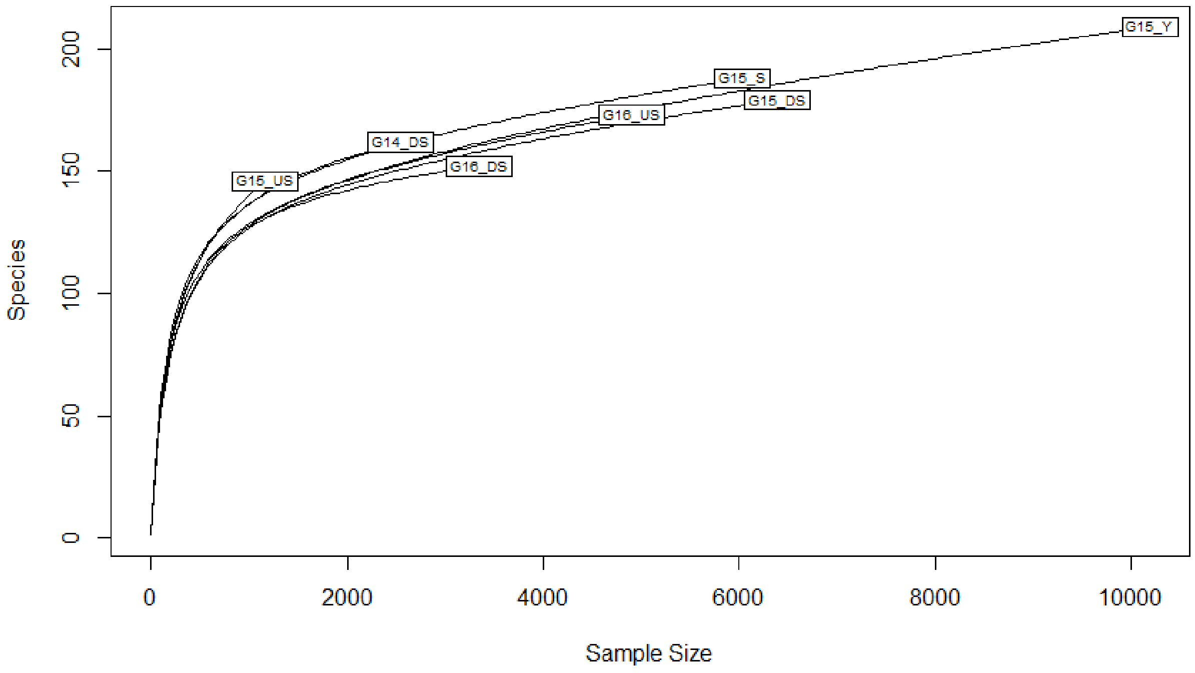


```
> setwd("E:/vinay/fungal_paper_data/")
>
> #Loading biom table
> slashpile_16sV1V3 <- "merge.biom"
> s16sV1V3 = import_biom(BIOMfilename = slashpile_16sV1V3, parseFunction = parse_taxonomy_default)
> colnames(tax_table(s16sV1V3)) <- c("Kingdom", "Phylum", "Class", "Order", "Family", "Genus")
> head(otu_table(s16sV1V3))
OTU Table:          [6 taxa and 7 samples]
                     taxa are rows
       G14_DS G15_US G15_Y G15_S G15_DS G16_US G16_DS
OTU263      0      0     0     0      0      0      0
OTU228      9      1    47    28     25      9     17
OTU260      6      3    24    14      7      9      8
OTU26      32     19   111    70     71     52     37
OTU98      14      1    47    28     21     13     14
OTU194     24      8   131    70     58     39     30
> head(sample_data(s16sV1V3))
Sample Data:        [6 samples by 3 sample variables]:
       Sample_name            Location              Considered_as
G14_DS      G14_DS    Kanpur_Downstream  Upstream_of_the_confluence
G15_US      G15_US   Allahabad_Upstream  Upstream_of_the_confluence
G15_Y        G15_Y     Allahabad_Yamuna                      Yamuna
G15_S        G15_S     Allahabad_Sangam                  Confluence
G15_DS      G15_DS Allahabad_Downstream Downstream_of_the_confluence
G16_US      G16_US    Varanasi_Upstream Downstream_of_the_confluence
> summarize_phyloseq(s16sV1V3)
Compositional = NO
```

```
1] Min. number of reads = 199
2] Max. number of reads = 1890
3] Total number of reads = 6224
4] Average number of reads = 889.142857142857
5] Median number of reads = 803
7] Sparsity = 0.213114754098361
6] Any OTU sum to 1 or less? YES
8] Number of singletons = 13
9] Percent of OTUs that are singletons 21.3114754098361
10] Number of sample variables are: 3
Sample_name
Location
Considered_as
>
> #Set Variable
> ps1 <- s16sV1V3
> summarize_phyloseq(ps1)
Compositional = NO
1] Min. number of reads = 199
2] Max. number of reads = 1890
3] Total number of reads = 6224
4] Average number of reads = 889.142857142857
5] Median number of reads = 803
7] Sparsity = 0.213114754098361
6] Any OTU sum to 1 or less? YES
8] Number of singletons = 13
9] Percent of OTUs that are singletons 21.3114754098361
10] Number of sample variables are: 3
Sample_name
Location
Considered_as
> print(ps1)
phyloseq-class experiment-level object
otu_table()   OTU Table:         [ 61 taxa and 7 samples ]
sample_data() Sample Data:       [ 7 samples by 3 sample variables ]
tax_table()   Taxonomy Table:    [ 61 taxa by 6 taxonomic ranks ]
>
> #Check if any OTUs are not present in any sample
> any(taxa_sums(ps1) == 0)
[1] TRUE
>
> #Removing of OTUs not present in any of the sample
> ps1a <- prune_taxa(taxa_sums(ps1) > 0, ps1)
> any(taxa_sums(ps1a) == 0)
[1] FALSE
> ntaxa(ps1)
[1] 61
> ntaxa(ps1a)
[1] 55
> ntaxa(ps1) - ntaxa(ps1a)
[1] 6
> rank_names(ps1a)
[1] "Kingdom" "Phylum"  "Class"   "Order"   "Family"  "Genus"
>
> #Plot Sequencing depth of each sample
```

```
> SeqDepth = colSums(otu_table(ps1a))
> sample_data(ps1a)$SeqDepth = SeqDepth
> ggplot(meta(ps1a)) +
+   geom_histogram(aes(x = log10(SeqDepth)), alpha= 0.6) + facet_wrap(~Sample_name) + theme_bw()
`stat_bin()` using `bins = 30`. Pick better value with `binwidth`.
> sort(SeqDepth)
G15_US G14_DS G16_DS G16_US G15_DS  G15_S  G15_Y
   199    457    578    803   1105   1192   1890
> min(SeqDepth)
[1] 199
> max(SeqDepth)
[1] 1890
> head(meta(ps1a))
       Sample_name            Location               Considered_as
G14_DS      G14_DS    Kanpur_Downstream   Upstream_of_the_confluence
G15_US      G15_US   Allahabad_Upstream   Upstream_of_the_confluence
G15_Y        G15_Y     Allahabad_Yamuna                       Yamuna
G15_S        G15_S     Allahabad_Sangam                   Confluence
G15_DS      G15_DS Allahabad_Downstream Downstream_of_the_confluence
G16_US      G16_US    Varanasi_Upstream Downstream_of_the_confluence
       SeqDepth
G14_DS      457
G15_US      199
G15_Y      1890
G15_S      1192
G15_DS     1105
G16_US      803
> ggbarplot(meta(ps1a), "Sample_name", "SeqDepth", fill = "Sample_name") + rotate_x_text()
> summarize_phyloseq(ps1a)
Compositional = NO
1] Min. number of reads = 199
2] Max. number of reads = 1890
3] Total number of reads = 6224
4] Average number of reads = 889.142857142857
5] Median number of reads = 803
7] Sparsity = 0.127272727272727
6] Any OTU sum to 1 or less? YES
8] Number of singletons = 7
9] Percent of OTUs that are singletons 12.7272727272727
10] Number of sample variables are: 4
Sample_name
Location
Considered_as
SeqDepth
>
> #Remove_singletons
> ps1b <- prune_taxa(taxa_sums(ps1a) > 1, ps1a)
> ps1b
phyloseq-class experiment-level object
otu_table()   OTU Table:         [ 48 taxa and 7 samples ]
sample_data() Sample Data:       [ 7 samples by 4 sample variables ]
tax_table()   Taxonomy Table:    [ 48 taxa by 6 taxonomic ranks ]
> summarize_phyloseq(ps1b)
Compositional = NO
1] Min. number of reads = 199
```

```
2] Max. number of reads = 1887
3] Total number of reads = 6217
4] Average number of reads = 888.142857142857
5] Median number of reads = 803
7] Sparsity = 0.0208333333333333
6] Any OTU sum to 1 or less? NO
8] Number of singletons = 0
9] Percent of OTUs that are singletons 0
10] Number of sample variables are: 4
Sample_name
Location
Considered_as
SeqDepth
>
> #otu_ditribution
> hist(log10(taxa_sums(ps1b)))
>
>
> #alpha_diversity
> alpha_meas = c("Shannon", "Simpson")
> (p <- plot_richness(ps1a, "Sample_name", "Sample_name", measures=alpha_meas))
> p + geom_boxplot(data=p$data, aes(x=Considered_as, y=value, color=NULL), alpha=0.1)
>
> #beta_diversity
> nsamples(ps1b)
[1] 7
>
> #set prevalence (we keept only those OTUs that are detected alteast 2 times in 2 out of total 7 samples)
> ps4 <- core(ps1b, detection = 2, prevalence = 2/nsamples(ps1b))
> ps4
phyloseq-class experiment-level object
otu_table()   OTU Table:         [ 47 taxa and 7 samples ]
sample_data() Sample Data:       [ 7 samples by 4 sample variables ]
tax_table()   Taxonomy Table:    [ 47 taxa by 6 taxonomic ranks ]
> ps4.rel <- microbiome::transform(ps4, "compositional")
> bx.ord_pcoa_bray <- ordinate(ps4.rel, "PCoA", "bray")
> beta.ps1 <- plot_ordination(ps4.rel,
+                             bx.ord_pcoa_bray,
+                             color="Sample_name",
+                             label = "Sample_name") +
+   geom_point(aes(shape = Sample_name), size= 4) +
+   theme(plot.title = element_text(hjust = 0, size = 12))
> beta.ps1 <- beta.ps1 + theme_bw(base_size = 14) +
+   theme(panel.grid.major = element_blank(), panel.grid.minor = element_blank())
> beta.ps2 <- beta.ps1 + geom_line() + scale_color_brewer(palette = "Dark2")
> beta.ps2
geom_path: Each group consists of only one observation. Do you need to
adjust the group aesthetic?
Warning messages:
1: The shape palette can deal with a maximum of 6 discrete values
because more than 6 becomes difficult to discriminate; you have 7.
Consider specifying shapes manually if you must have them.
2: Removed 1 rows containing missing values (geom_point).
>
>
```

```
> ps1a.meta <- meta(ps1a)
> ps1a.meta$simpson <- ps.even$simpson
> hist(ps1a.meta$simpson)
>
> #shapiro test is used to check the distribution of otus ....
> shapiro.test(ps1a.meta$simpson)

        Shapiro-Wilk normality test

data:  ps1a.meta$simpson
W = 0.80444, p-value = 0.04531

> qqnorm(ps1a.meta$simpson)
> #Rarefaction_Plot
> rarecurve(t(otu_table(ps1a)), step=50, cex=0.5)
> #alpha_diversity
> alpha_meas = c("Simpson")
> (p <- plot_richness(ps1a, "Sample_name", "Sample_name", measures=alpha_meas))
> p + geom_boxplot(data=p$data, aes(x=Considered_as, y=value, color=NULL), alpha=0.1)



>
```

```
library("phyloseq")

library("ggplot2")

library("gridExtra")

library("edgeR")

library("vegan")

library("microbiome")

library("knitr")

library("ggpubr")

library("reshape2")

library("RColorBrewer")

library("microbiomeutilities")

library("viridis")

library("tibble")


#Set_working_directory

setwd("E:/vinay/fungal_paper_data/")
```

```
#Loading biom table
slashpile_16sV1V3 <- "merge.biom"
s16sV1V3 = import_biom(BIOMfilename = slashpile_16sV1V3, parseFunction =
parse_taxonomy_default)
colnames(tax_table(s16sV1V3)) <- c("Kingdom", "Phylum", "Class", "Order", "Family", "Genus")
head(otu_table(s16sV1V3))
head(sample_data(s16sV1V3))
summarize_phyloseq(s16sV1V3)

#Set Variable
ps1 <- s16sV1V3
summarize_phyloseq(ps1)
print(ps1)

#Check if any OTUs are not present in any sample
any(taxa_sums(ps1) == 0)

#Removing of OTUs not present in any of the sample
ps1a <- prune_taxa(taxa_sums(ps1) > 0, ps1)
any(taxa_sums(ps1a) == 0)
ntaxa(ps1)
ntaxa(ps1a)
ntaxa(ps1) - ntaxa(ps1a)
rank_names(ps1a)

#Plot Sequencing depth of each sample
SeqDepth = colSums(otu_table(ps1a))
sample_data(ps1a)$SeqDepth = SeqDepth
ggplot(meta(ps1a)) +
```

```
  geom_histogram(aes(x = log10(SeqDepth)), alpha= 0.6) + facet_wrap(~Sample_name) + theme_bw()
sort(SeqDepth)
min(SeqDepth)
max(SeqDepth)
head(meta(ps1a))
ggbarplot(meta(ps1a), "Sample_name", "SeqDepth", fill = "Sample_name") + rotate_x_text()
summarize_phyloseq(ps1a)

#Remove_singletons
ps1b <- prune_taxa(taxa_sums(ps1a) > 1, ps1a)
ps1b
summarize_phyloseq(ps1b)

#otu_ditribution
hist(log10(taxa_sums(ps1b)))



#alpha_diversity
alpha_meas = c("Simpson")
(p <- plot_richness(ps1a, "Sample_name", "Sample_name", measures=alpha_meas))
p + geom_boxplot(data=p$data, aes(x=Considered_as, y=value, color=NULL), alpha=0.1)

#beta_diversity
nsamples(ps1b)

#set prevalence (we keept only those OTUs that are detected alteast 2 times in 2 out of total 7 samples)
ps4 <- core(ps1b, detection = 2, prevalence = 2/nsamples(ps1b))
ps4
ps4.rel <- microbiome::transform(ps4, "compositional")
bx.ord_pcoa_bray <- ordinate(ps4.rel, "PCoA", "bray")
```

```
beta.ps1 <- plot_ordination(ps4.rel,
                            bx.ord_pcoa_bray,
                            color="Sample_name",
                            label = "Sample_name") +
  geom_point(aes(shape = Sample_name), size= 4) +
  theme(plot.title = element_text(hjust = 0, size = 12))
beta.ps1 <- beta.ps1 + theme_bw(base_size = 14) +
  theme(panel.grid.major = element_blank(), panel.grid.minor = element_blank())
beta.ps2 <- beta.ps1 + geom_line() + scale_color_brewer(palette = "Dark2")
beta.ps2


ps1a.meta <- meta(ps1a)
ps1a.meta$simpson <- ps.even$simpson
hist(ps1a.meta$simpson)

#shapiro test is used to check the distribution of otus ....
shapiro.test(ps1a.meta$simpson)
qqnorm(ps1a.meta$simpson)

#taxonomy_composition
>ps1a.com <- ps1a
>taxic <- as.data.frame(ps1a.com@tax_table)
>taxic$OTU <- rownames(taxic)
>colnames(taxic)
>taxmat <- as.matrix(taxic)
>new.tax <- tax_table(taxmat)
>tax_table(ps1a.com) <- new.tax
>pseq.ph <- aggregate_taxa(ps1a.com, "Phylum", top = 11)
>p.phy <- plot_composition(pseq.ph, sample.sort = NULL, otu.sort = NULL,
```

```
                    x.label = "Description", plot.type = "barplot", verbose = FALSE)

>print(p.phy + scale_fill_brewer(palette = "Paired") + theme_bw())

#Relative_abundance
>guide_italics <- guides(fill = guide_legend(label.theme = element_text(size = 15,
                                                    face = "italic", colour = "Black", angle = 0)))
>pseq.ph.rel <- microbiome::transform(pseq.ph, "compositional")
>plot.comp.rel <- plot_composition(pseq.ph.rel, x.label = "SampleType") +
  theme(legend.position = "bottom") + theme_bw() +
  theme(axis.text.x = element_text(angle = 90)) +
  ggtitle("Relative abundance") + guide_italics +
  theme(legend.title = element_text(size=18))

plot.comp.rel + scale_fill_brewer( "Phylum",palette = "Paired")

#Rarefaction_Plot
rarecurve(t(otu_table(ps1a)), step=50, cex=0.5)
```

**Supplementary S5.** OUT distribution scatter plot

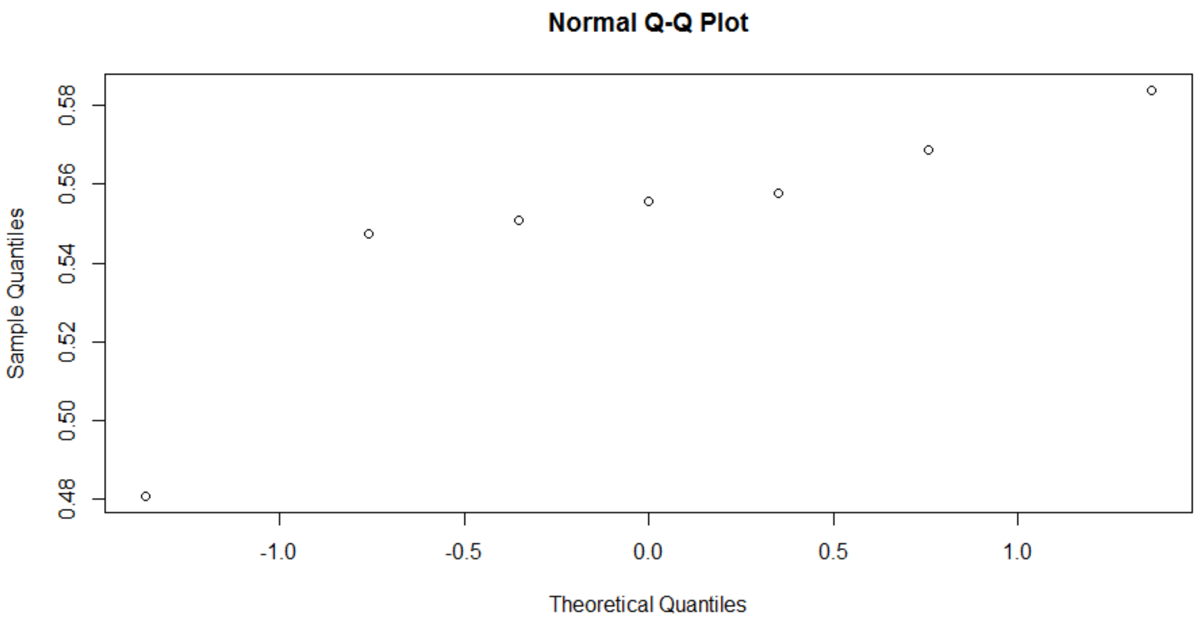
